# Arginine induced *Streptococcus gordonii* biofilm detachment using a novel rotating-disc rheometry method

**DOI:** 10.1101/2021.03.15.435500

**Authors:** Erin S. Gloag, Daniel J. Wozniak, Kevin L. Wolf, James G. Masters, Carlo Amorin Daep, Paul Stoodley

**Affiliations:** Department of Microbial Infection and Immunity, The Ohio State University, Columbus, OH, USA, 43210; Department of Microbiology, The Ohio State University, Columbus, OH, USA, 43210; Department of Mechanical and Aerospace Engineering, The Ohio State University, Columbus, Ohio, USA, 43210; Colgate-Palmolive Technology Center, Piscataway, NJ, USA, 08854; Department of Orthopedics, The Ohio State University, Columbus, OH, USA, 43210; National Biofilm Innovation Centre (NBIC) and National Centre for Advanced Tribology at Southampton (nCATS), University of Southampton, Southampton, UK, SO17 1BJ

**Author notes:** Correspondence to: Erin S. Gloag, 760 BRT, 460W 12^th^ Ave Columbus OH, 43210, USA.

**Keywords:** viscoelasticity, biophysical, mechanics, *Streptococcus gordonii*, arginine, dental plaque

## Abstract

Oral diseases are one of the most common pathologies affecting human health. These diseases are typically associated with dental plaque-biofilms, through either build-up of the biofilm or dysbiosis of the microbial community. Arginine can disrupt dental plaque-biofilms, and maintain plaque homeostasis, making it an ideal therapeutic to combat the development of oral disease. Despite our understanding of the actions of arginine towards dental plaque-biofilms, it is still unclear how or if arginine effects the mechanical integrity of the dental plaque-biofilm. Here we adapted a rotating-disc rheometry assay, which is routinely used in marine microbial ecology, to study how arginine treatment of *Streptococcus gordonii* biofilms influences biofilm detachment from surfaces. We demonstrate that the assay is highly sensitive at quantifying the presence of biofilm and the detachment or rearrangement of the biofilm structure as a function of shear stress. We demonstrate that arginine treatment leads to earlier detachment of the biofilm, indicating that arginine treatment weakens the biofilm, making it more suspectable to removal by shear stresses. Our results add to the understanding that arginine targets biofilms by multifaceted mechanisms, both metabolic and physical, further promoting the potential of arginine as an active compound in dentifrices to maintain oral health.

## Introduction

Biofilms are communities of microorganisms, encased in an extracellular polymeric slime (EPS). These communities adhere at either surface interfaces or to neighboring microorganisms (Bjarnsholt *et al,* 2013). Biofilms are responsible for a number of infectious diseases, where these communities are highly recalcitrant to traditional therapies, promoting the persistence of these infections (Hall-Stoodley *et al,* 2004). Dental plaque is perhaps one of the most widely understood biofilms affecting human health. Oral pathologies typically arise due to poor oral hygiene and diet, that lead to dental plaque build-up or dysbiosis of the plaque microbial community. Together these factors can lead to oral diseases including dental caries, gingivitis and periodontitis (Mosaddad *et al,* 2019). Oral hygiene, including combinations of mechanical dental plaque removal and antimicrobial agents in dentifrices, continues to be the most effective method at preventing the development of these pathologies.

Exogenous arginine has emerged as a novel therapy to combat dental plaque. This mechanism has been chiefly attributed to the buffering capacity of arginine metabolism by arginolytic organisms, including *Streptococcus gordonii*. These organisms encode an arginine deiminase system (ADS), which metabolizes arginine, producing ammonia (Jakubovics *et al,* 2015, Wijeyeweera *et al,* 1989a, b). This in turn neutralizes acid produced by acidogenic organisms, maintaining a neutral pH within the dental plaque-biofilm (Wijeyeweera *et al,* 1989a, b). Exogenous arginine treatment also promotes *S. gordonii* growth and prevents the out-growth of cariogenic species, including *Streptococcus mutans,* in mixed species biofilm models (Bijle *et al,* 2019, He *et al,* 2016).

Exogenous arginine treatment can also reduce microbial coaggregation (Ellen *et al,* 1992, Kamaguch *et al,* 2001, Levesque *et al,* 2003), and alters the EPS biochemical composition, by preventing the out-growth of *S. mutans*, and subsequently reducing the amount of insoluble glycans produced by this organism (He *et al,* 2016, Kolderman *et al,* 2015). Interestingly, treatment with low concentrations of arginine promotes the growth of *S. gordonii* biofilms, however, high concentrations of the amino acid reduces biofilm biomass (Jakubovics *et al,* 2015). It was predicted that arginine treatment inhibited cell-cell interactions within the biofilm (Jakubovics *et al,* 2015). Taken together these data suggest that exogenous arginine treatment can disrupt dental plaque-biofilm, preventing its build-up (Wolff *et al,* 2018, Kolderman *et al,* 2015, Manus *et al,* 2018).

Despite the above observations, there is little understanding of how arginine treatment impacts the mechanical integrity of dental plaque-biofilms, an important factor in understanding how antimicrobials may penetrate the biofilm or how mechanical disruption may physically remove the biofilm. Atomic force microscopy (AFM) showed that *S. mutans* biofilms, grown in the presence of arginine, had reduced adhesion forces to the AFM tip (Sharma *et al,* 2014). This was predicted to be due to reduced glycan production or hydrogen bonds within the EPS (Sharma *et al,* 2014). However, effects of arginine treatment on the bulk biofilm properties and biofilm removal have yet to be considered. Furthermore, most studies have focused on how arginine impacts *S. mutans* biofilms, or caries-active plaque (Wolff *et al,* 2018). Few have focused on understanding how arginine impacts non-cariogenic plaque, or the biofilms of early plaque colonizers, such as *S. gordonii* (Jakubovics et al, 2015).

Rotating discs have long been used to analyze how biofilm fouling effects the hydrodynamics and drag associated with marine biofouling (Granville, 1982). The disc is rotated at increasing angular velocity, and the resulting torque (resistance to imparted rotary motion) is measured. Increases in torque is related to biomass, roughness and deformability of the biofilm (Dennington *et al,* 2015). Conventionally, such discs are large (i.e. between 0. 5 - 1 m diameter), and hence cumbersome to manage. However, recently non-contact rotating-disc rheometry has been used to analyze drag associated with marine biofouling on discs 2.5 - 4 cm in diameter (Dennington *et al,* 2015). In this method a rheometer is used as a highly sensitive torque monitor, allowing precise measurements of torque, even that generated by small discs compatible with the scale of routine laboratory biofilm growth systems (Dennington *et al,* 2015). As such, it represents a novel method for direct quantification of biofilms outside of traditional assays, such as microscopic examination, viable counts and crystal violet staining. In addition, it allows real time correlation between imposed shear stress and real time changes in torque when biofilm is detached, informing how much shear is required to disrupt the biofilm. Here we adapted rotating-disc rheometry to study *S. gordonii* biofilm detachment after arginine treatment.

## Materials and Methods

### 3D printing coupons

The model for the coupons was designed in SolidWorks (Dassault Systèmes); refer to supplementary file 1. Coupons were 3D printed using a Prime 30 PolyJet 3D printer (Objet, Stratasys) using RGD720 photopolymer for the printing material (Stratasys). The coupon was printed at a resolution of 0.0008 inches. The coupon surface was sanded used P300 sandpaper to create a rougher surface for bacteria to attach. Prior to inoculating, coupons were sterilized in 70% ethanol.

### *S. gordonii* biofilm growth and treatment

*S. gordonii* wild type strain DL1 was used in this study. Overnight cultures were prepared by inoculating 10 mL of brain heart infusion broth (Oxoid; BHI) with a colony of *S. gordonii* and incubated statically overnight at 37°C with 5% CO_2_.

Sterile 40 mm coupons were placed in a Petri dish containing 40 mL BHI, supplemented with 0.5% sucrose. Coupons were inoculated with 400 μL of overnight culture. Biofilms were incubated in a humidified chamber at 37°C with 5% CO_2_, on an orbital shaker at 150 rpm. Every 24 h the media was replenished. Biofilms were grown for 5 days.

Biofilms were treated by transferring the coupons to a Petri dish containing either 40 mL PBS or 4% arginine. Biofilms were treated for 2 min at 37°C with 5% CO_2_, shaking at 150 rpm. Biofilms were washed in PBS and transferred to 40 mL PBS until analysis.

### Adapted rotating-disc rheometry analysis

Biofilms were analyzed on a Discovery Hybrid Rheometer-2 (HD-2) (TA Instruments). A 15 × 15 cm square clear acrylic container filled with 2.8 L reverse osmosis water was transferred onto the Peltier plate. Biofilm-coated coupons were immersed and attached to the rheometer shaft using a custom-made adapter probe. The gap distance between the bottom of the container and the coupon was set to 3.5 cm (Fig 1A). Immersed coupons were spun at an angular velocity (ω) range of 0.1 – 300 rad·s^−1^, incrementing the speed across 360 s. Four biological replicates were analyzed, each with 2 – 4 technical replicates (total N = 11). It is important to note that the geometry of the system will influence the motion of water in the reservoir. As such measurements should be considered system-specific.

**Figure 1:**
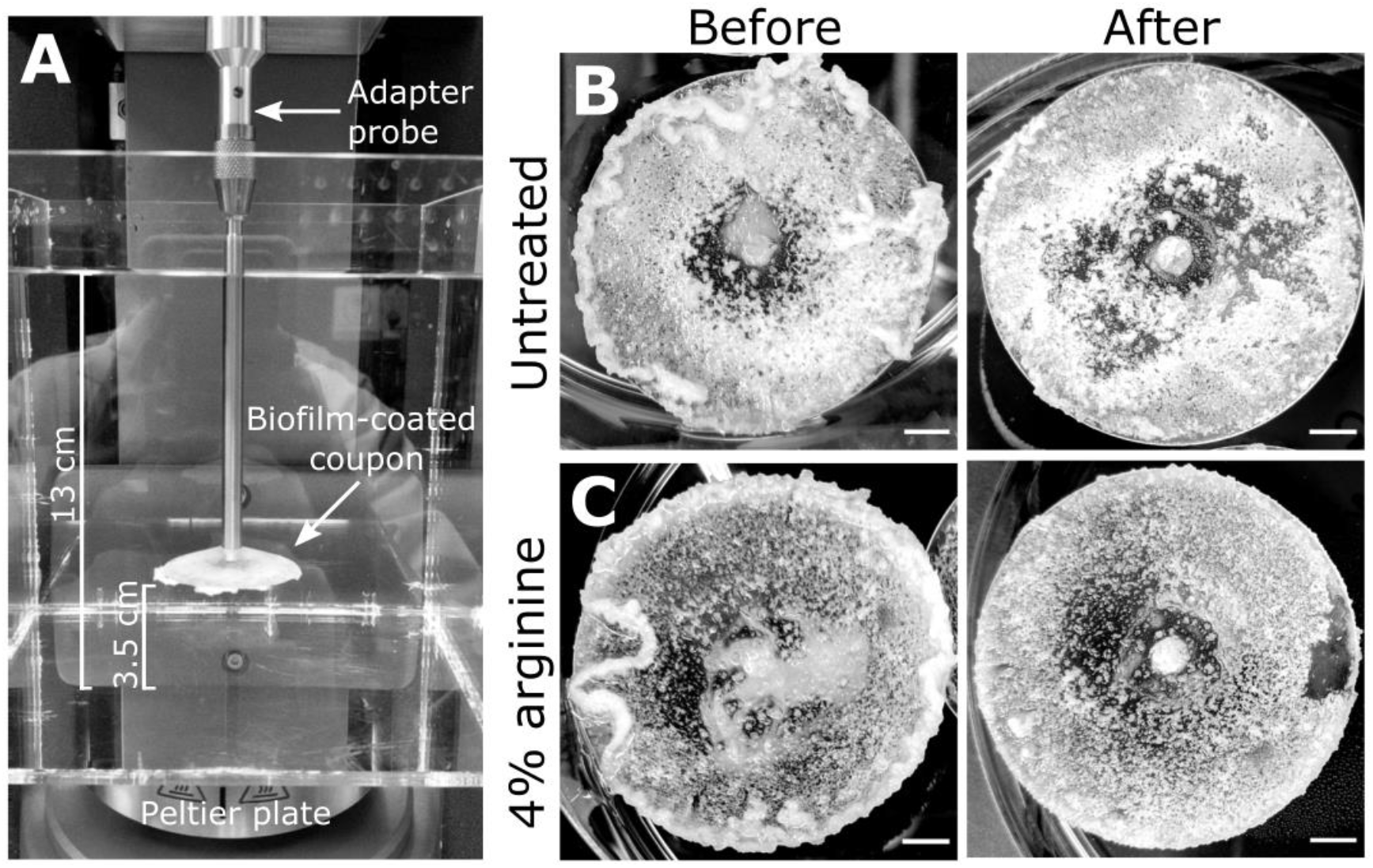
*S. gordonii* biofilms before and after analysis. **(A)** Experimental design for the adapted rotating-disc rheometry analysis. Biofilm-coated coupons were attached to an adaptor probe on the rheometer using a threaded tap that was printed onto the back of the coupon. This was immersed in a container filled with reverse osmosis water. A gap thickness of 3.5 cm was set between the coupon and the bottom of the container. Prior to analysis 5 day *S. gordonii* biofilms, grown on the coupons, were treated with either **(B)** PBS (untreated control) or with **(C)** 4% arginine (labeled). Images depict biofilms before and after rheometry analysis (labeled). Scale bar indicates 5mm.

### Data Analysis

Data was collected using TRIOS v5 software (TA instruments), with raw data exported in excel. Data was transformed, and calculations performed in excel. Data was visualized and statistical analysis performed in GraphPad Prism v8 (GraphPad Software). All statistical comparisons were performed using a Student’s *t*-test, with *p* < 0.05 indicating significance.

To more clearly observe the changes in torque, the torque – angular velocity curves were linearized and transformed (Fig S1). The data was linearized by taking the square root of the torque. The running slope of 5 data points of the linearized data was determined. This transformed data was linearized after 20 rad·s^−1^. Therefore, final transformed data is presented as the running slope of the linearized data against angular velocity, starting at 20 rad·s^−1^ (Fig S1).

The biofilm momentum coefficient *(C_B_)*, also referred to as the momentum or torque coefficient, was determined as previously described (Dennington *et al,* 2015). The adapted rotating-disc rheology measurement is most sensitive at detecting changes in torque at the turbulent regime, between 200 – 300 rad·s^−1^. Torque within this range has a linear relationship to ω^2^, where the slope of this line *(T^1/2^/ω)* equates to *C_B_ · k*. Therefore, *C_B_* can be defined by equation 1:

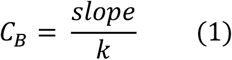

where *k* is a constant for the system, defined by:

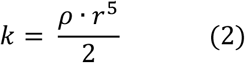

where ρ is the density of the fluid, in this case water (997 kg/m^3^) and r is the radius of the coupon (0.02 m).

The angular velocity where the first decrease in torque was detected was converted to the shear stress acting at the outer edge of the coupon (τ), as previously described (Hunsucker *et al,* 2016), according to equation 3:

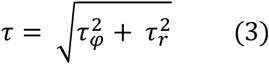

where, τ_φ_ is the shear stress acting in the circumferential direction and τ_r_ is the shear stress acting radially.

The shear stress acting in the circumferential direction is described by equation 4:

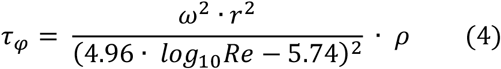

where Re is the Reynolds number acting at the outer edge of the coupon described by equation 5:

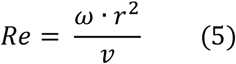

where v is the kinematic viscosity (9 × 10^−7^ m^2^·s^−1^).

The shear stress acting in the radial direction is described by equation 6:

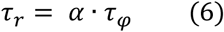

where α is the skewness between the shear stress acting in both directions, and is described by equation 7:

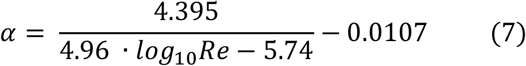

Finally, the area under the curve (AUC) of the torque – angular velocity curves was determined using the analysis function in GraphPad Prism.

## Results

### Adapted rotating-disc rheometry is sensitive at detecting biofilm rearrangement and detachment events

Mechanical analysis of biofilms is becoming more widespread in the field (Gloag *et al,* 2019). However, there is currently a lack in analyzing biofilm mechanics in the context of biofilm removal. To meet this need we adapted rotating-disc rheology to analyze biofilm detachment from surfaces.

*S. gordonii* biofilms were grown on 3D printed coupons for 5 days. Biofilm coated coupons were connected to the rheometer and immersed in reverse osmosis water (Fig 1A). Coupons were spun across an angular velocity range of 0.1 – 300 rad·s^−1^ over 360 s, and the resulting torque, a measurement of resistance to rotation, was measured (Movie S1). Across this velocity range, detachment of biofilm aggregates was observed, particularly at the higher velocity regimes. These detachment events appeared to correlate to reductions in torque (Movie S1), with both small (Fig 2A) and larger (Fig 2B) aggregate detachments detected. After analysis there remained biofilm still attached to the coupon surface (Fig 1B). The remaining biofilm was not removed with repeated analysis (Fig S2).

**Figure 2:**
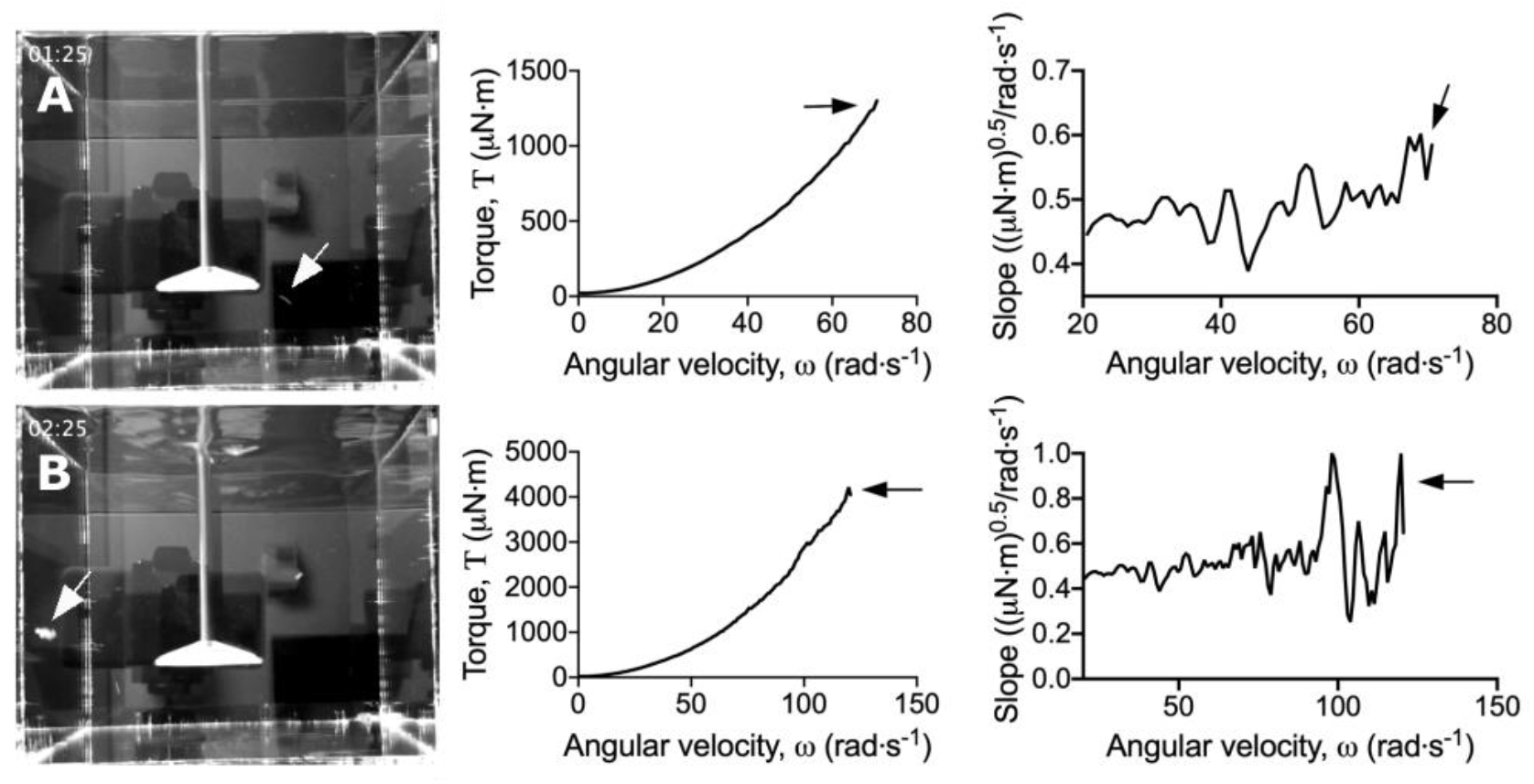
Dips in the torque – angular velocity curve correlate to biofilm detachment events from the coupon. Stills taken from movies S1 and S2 (left panel) depicting **(A)** small and **(B)** large biofilm detachment events. The torque – angular velocity curve (middle panel) and transformed linearized analysis (right panel) at each time is depicted. White arrows indicate detached biofilm and black arrows indicate the corresponding changes in the curve.

To more easily observe the changes in torque, associated with biofilm detachment, the torque – angular velocity data was linearized and transformed (Fig S1). Using this transformed analysis, the reductions in torque were emphasized (Movie S2; Fig 2). Furthermore, changes in torque not associated with macroscopic aggregate detachment were observed, particularly at the lower velocity regimes (Movie S2). This suggested that the adapted rotating-disc rheometry analysis was capable of detecting microscopic detachment events, or rearrangement of the biofilm structure in response to external shear stress.

### 4% arginine treated biofilms are more sensitive to removal by shear stresses

Having validated the sensitivity of the adapted rotating-disc rheometry, we used this assay to determine how arginine treatment influenced biofilm mechanics, in regards to biofilm removal. Five day *S. gordonii* biofilms were treated with either PBS (untreated control) or 4% arginine for 2 min. This short treatment time was selected to mimic the time that a person would typically brush their teeth.

Macroscopically, arginine treatment did not appear to affect biofilm morphology, or the amount of remaining biofilm attached to the coupon after rheometry analysis (Fig 1B, C). However, arginine treated biofilms displayed reduced torque, compared to untreated biofilms (Fig 3). This indicates that coupons with arginine treated biofilms could rotate more easily across the assayed angular velocity range. This is further highlighted by the transformed data, where greater changes in torque, indicated by more negative slope values, were observed for arginine treated biofilms, compared to untreated (Fig 4). Both treated and untreated biofilms had increased torque values compared to the coupon alone (Fig 3 and 4).

**Figure 3:**
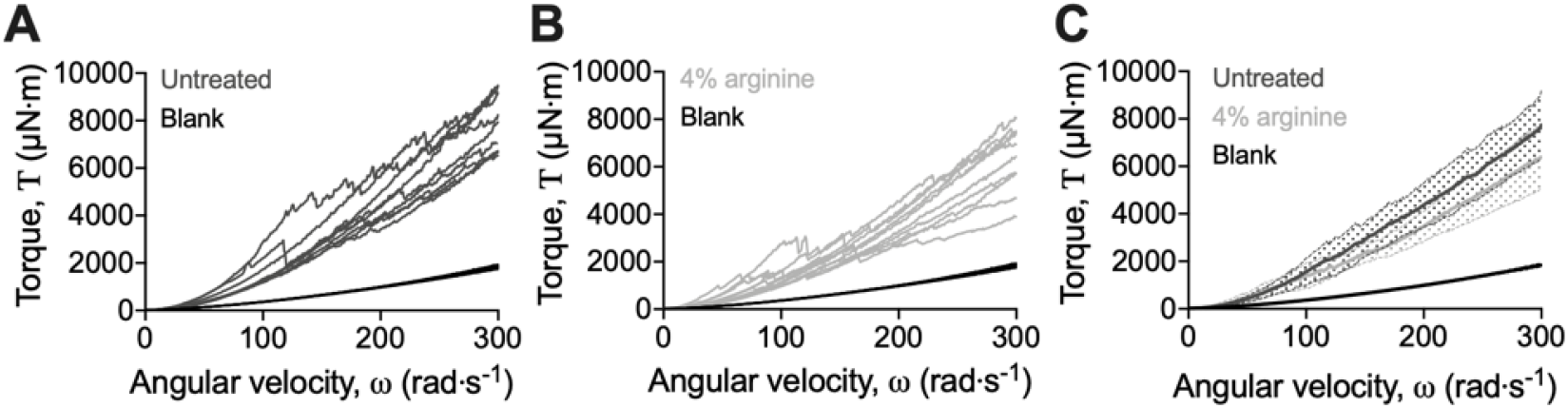
Adapted rotating-disc measurements of untreated and 4% arginine treated *S. gordonii* biofilms. Curves of individual **(A)** untreated and **(B)** 4% arginine treated *S. gordonii* biofilms. **(C)** Data from **(A)** and **(B)** expressed as mean ± standard deviation. In each panel, blank indicates analysis for coupon alone, with no biofilm. 4 biological replicates were performed, with 2 – 4 biofilms analyzed for each replicate (total N = 11).

**Figure 4:**
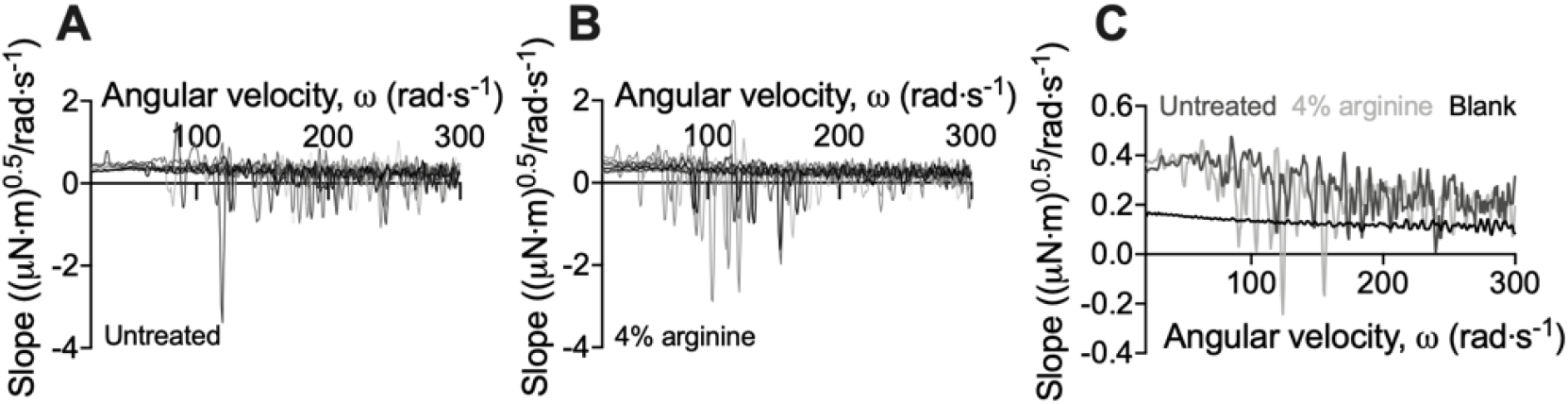
Transformed linearized analysis of untreated and 4% arginine treated *S. gordonii* biofilms. Curves of individual **(A)** untreated and **(B)** 4% arginine treated *S. gordonii* biofilms. **(C)** Data from **(A)** and **(B)** expressed as mean. In each panel, blank indicates analysis for coupon alone, with no biofilm. 4 biological replicates were performed, with 2 – 4 biofilms analyzed for each replicate (total N = 11).

To quantify the mechanical differences between arginine treated and untreated *S. gordonii* biofilms, the biofilm momentum coefficient across the turbulent regimes of 200 – 300 rad·s^−1^, was determined according the equ 1. The biofilm momentum coefficient is a dimensionless unit that is an indication of the drag caused by the biofilm, which in turn is related to the thickness and roughness of biofilm. Therefore, a higher coefficient is associated with more drag on the coupon, due to increased amount of adhered biofilm (Dennington *et al,* 2015, Granville, 1982). Both untreated and treated *S. gordonii* biofilms had biofilm momentum coefficients significantly greater than the coupon alone (Fig 5A), indicating that the presence of attached biofilm significantly hampered rotation of the coupon. Although there was no statistical difference between the coefficient of untreated and treated biofilms, there was greater variability for arginine treated biofilms, as indicated by the standard deviation; 0.006 for untreated biofilms and 0.013 for arginine treated biofilms (Fig 5A). This indicates that there was greater heterogeneity of arginine treated biofilms across 200 – 300 rad·s^−1^, compared to untreated biofilms. To look into these differences further, the AUC of the torque – angular velocity curves was determined (Fig 5B). Unlike the biofilm momentum coefficient, which only takes into consideration coupon rotation between 200 – 300 rad·s^−1^, AUC considers the rotation across the whole analyzed range. From this analysis, both untreated and treated *S. gordonii* biofilms had significantly greater AUC compared to the coupon alone, consistent with the fouling coefficient (Fig 5B). However, arginine treated biofilms had significantly reduced AUC, compared to untreated biofilms (Fig 5B). This suggests that, when also considering the lower velocity ranges, less work was required for rotation of the coupon, compared to untreated biofilms.

**Figure 5:**
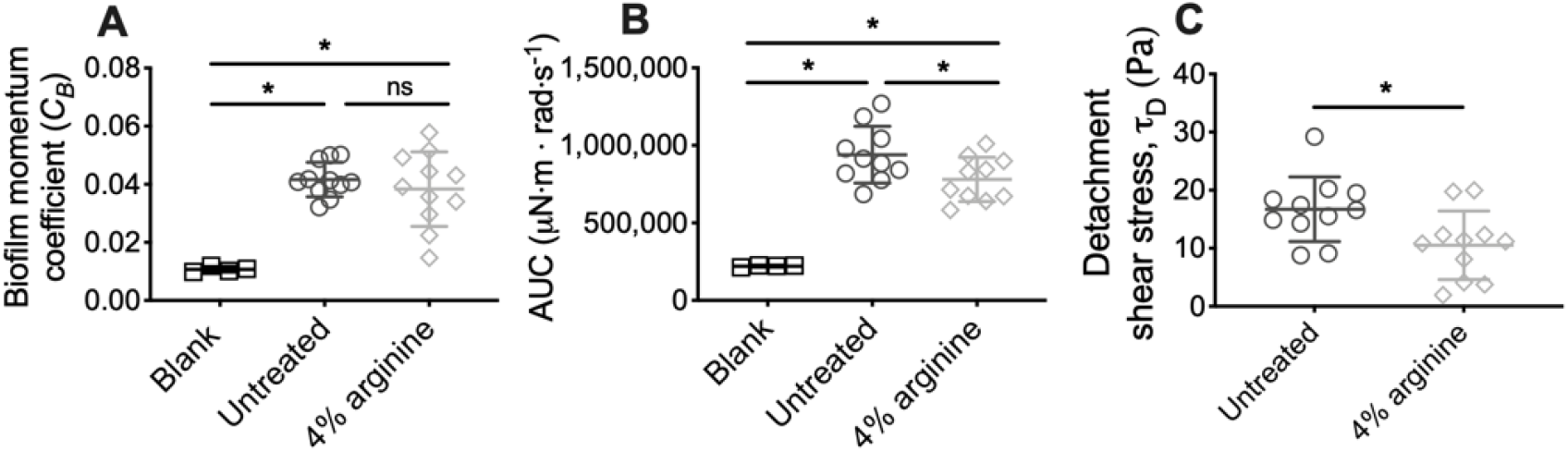
Treatment with 4% arginine weakens *S. gordonii* biofilms. **(A)** Biofilm momentum coefficient *(C_B_)*, determined according to equ 1, at 200 – 300 rad.s^−1^ in Fig 3. **(B)** Area under the curve (AUC) of torque – angular velocity curves depicted in Fig 3. **(C)** Initiation of detachment, indicated as the first reduction in torque in Fig 3, converted to shear stress according to equ 3. * p-value < 0.05, ns indicates no statistical difference.

From the transformed analysis, it also appeared that for arginine treated biofilms, the changes in torque, associated with biofilm detachment events, occurred at lower angular velocity ranges, compared to untreated biofilms (Fig 4). To quantify these differences, the angular velocity where the first reduction in torque occurred was converted to the shear stress acting on the outer edge of the coupon, according to equ 3. This analysis revealed that reductions in torque occurred at significantly lower shear stresses for arginine treated biofilms, compared to untreated (Fig 5C). This indicates that arginine treated biofilms were detaching from coupons at lower shear stresses, suggesting that they were more easily removed by external shear forces, compared to untreated biofilms.

## Discussion

Arginine is emerging as a potential therapeutic to prevent oral diseases, due to its ability to maintain dental plaque-biofilm homeostasis and disrupt biofilm formation (Wolff *et al,* 2018, Kolderman *et al,* 2015, Manus *et al,* 2018). However, there remains little understanding of how arginine treatment impacts biofilm mechanics or detachment. Here we adapted rotating-disc rheometry from the field of biofouling (Dennington *et al,* 2015, Granville, 1982), to study how shear induced removal of *S. gordonii* biofilms was effected by arginine treatment.

Our data suggests that *S. gordonii* biofilms appear to consist of two layers. An upper layer that was readily removed, and a base layer that was more adherent, and resistant to removal (Fig S2). This was true for both arginine treated and untreated *S. gordonii* biofilms (Fig 1B and C). Similarly, a remaining biofilm layer that was resistant to removal when exposed to increasing shear stresses was observed for *S. mutans* biofilms (Hwang *et al,* 2014), and biofilms grown from river (Desmond *et al,* 2018) and drinking (Abe *et al,* 2012) waters. Mechanical heterogeneity across the biofilm *z-*plane architecture has also been quantified for *Pseudomonas fluorescens* (Cao *et al,* 2016) and *Escherichia coli* (Galy *et al,* 2012) biofilms using micro-rheology methods. Together, this suggests that common to biofilms are a stratified mechanical architecture, resulting in a cohesion/ adhesion gradient, with the base of the biofilm being ridge and highly resistant to external forces. This could have important implications when considering the mechanical and chemical removal of biofilms from surfaces.

Our analysis also revealed that arginine treated *S. gordonii* biofilms had both reduced drag on the coupon during rotation (Fig 5A and B), and detached from the coupon at lower shear stresses (Fig 5C), compared to untreated biofilms. This suggests that arginine treatment weakened the structure of *S. gordonii* biofilms and that they were more easily removed from surfaces by external mechanical forces. Interestingly, previous observations of the biofilm disrupting effects of arginine either grew the biofilms in the presence of arginine, or treated the biofilms at multiple time points (He *et al*, 2016, Kolderman *et al*, 2015, Jakubovics *et al*, 2015). When mixed species biofilms were treated with arginine, three times a day over approximately 2 days, arginine effects to both microbial populations and biofilm structure were observed after 53 h (He *et al,* 2016). It was determined that arginine treatment takes time to exert effects on the biofilm, suggesting that arginine metabolism by arginolytic bacteria is required (He *et al,* 2016). However, here we observed arginine weakening *S. gordonii* biofilms after only 2 min of treatment. This suggests that mechanical destabilization of the biofilm can occur within a rapid time frame, compared to those that visually impact the biofilm architecture. These immediate mechanical effects are likely due to physiological interactions, rather than metabolic.

AFM analysis of *S. mutans* biofilms, grown in the presence of arginine, identified that arginine reduced biofilm adhesion. *S. mutans* cannot metabolize arginine, and it was predicted that arginine prevented hydrogen bond interactions across glycan polymers within the EPS (Sharma *et al,* 2014). Furthermore, disruption of *S. gordonii* biofilms, when grown in the presence of high arginine concentrations, was predicted to be independent of arginine metabolism. Rather, it was predicted to be due to inhibition of cell-cell interactions within the biofilm (Jakubovics *et al,* 2015). We therefore predict that the weakening of arginine treated *S. gordonii* biofilms observed here, may be due to disruption of chemical interactions between EPS components, or cell-cell or cell-EPS interactions within the biofilm. Similarly, *S. mutans* biofilms treated with a hydrolase that degrades EPS, were more easily removed from surfaces by exposure to external shear forces (Hwang *et al,* 2014).

Here we have adapted rotating-disc rheometry from the field of biofouling, as a novel methodology to analyze biofilm detachment from surfaces. We demonstrated that this assay is highly sensitive at detecting biofilm detachment, and possible structural rearrangements, with increasing shear forces. This methodology is also sensitive at detecting mechanical changes to the biofilm architecture that are not visually apparent. However, this method is destructive to the biofilm, and therefore, limits the sensitivity of assessing drag of the original structure. Finally, we also identified, for the first time, that arginine treatment can weaken the mechanical structure of *S. gordonii* biofilms, resulting in detachment at lower shear stresses, compared to untreated biofilms. These effects were observed after only 2 min of treatment. These results add to the multifaceted action of arginine at disrupting dental plaque-biofilms, and further promotes the potential use of arginine as an active compound in dentifrices to combat dental plaque and help improve oral health.

## Acknowledgements

This work was funded by Colgate-Palmolive. ESG was funded by an American Heart Association Career Development Award (19CDA34630005). DJW and PS were funded by the National Institute of Health (R01AI134895 and R01AI143916; DJW) (R01GM124436; PS).

## Author contributions

KLW designed and printed the coupons. ESG performed all experimental work. ESG and PS analyzed and interpreted the experimental data. ESG, DJW, CD and PS wrote the manuscript. All authors gave their final approval and agree to be accountable for all aspects of the work.

---

**Supplementary Figure 1:**
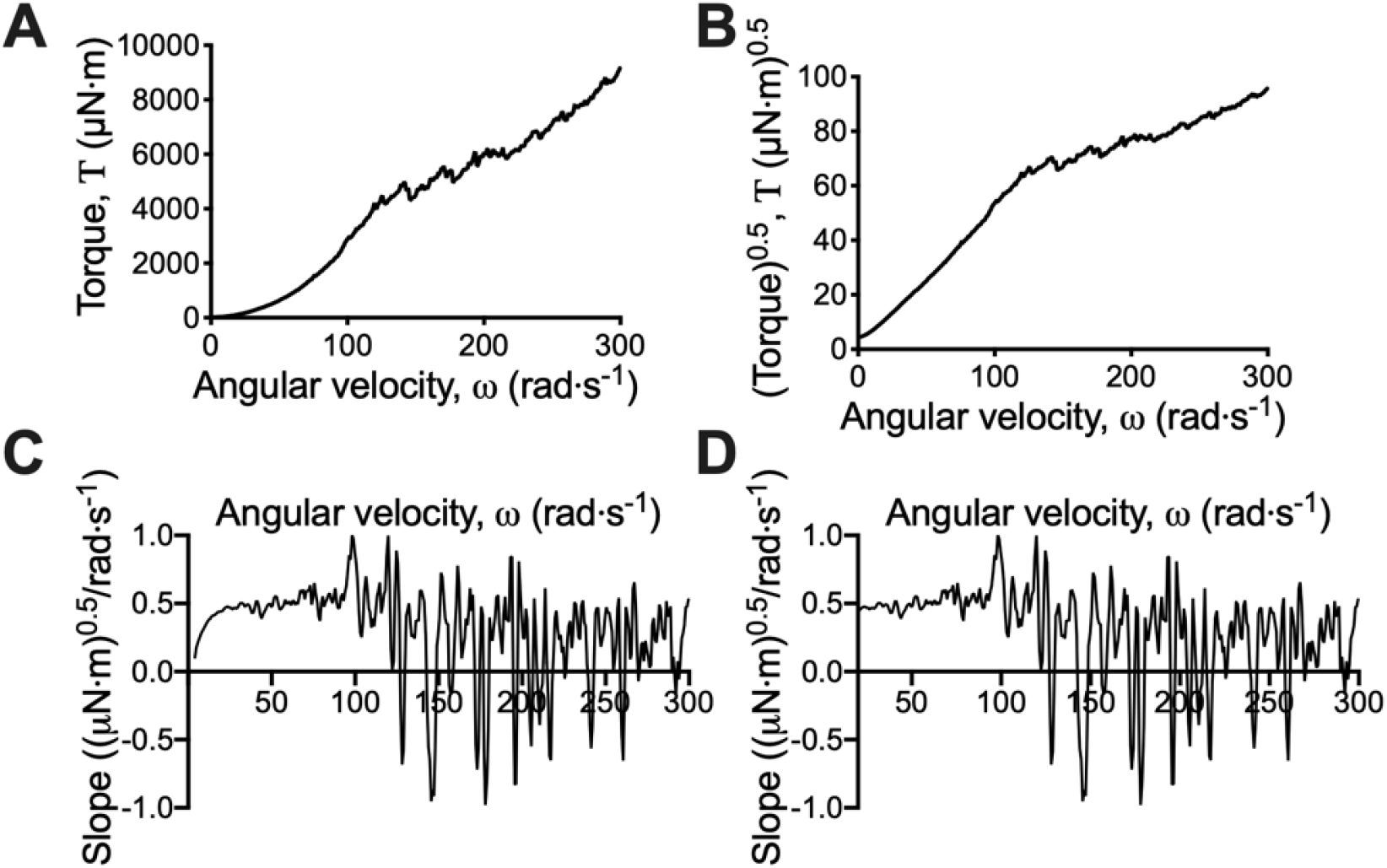
Data processing. **(A)** The torque and angular velocity data were exported from TRIOS v5 software. To visualize the changes in torque with angular velocity more clearly the data was linearized by **(B)** plotting the square root of the torque against angular velocity. **(C)** The running slope of 5 data points of the linearized data was plotted against the angular velocity. **(D)** This transformed data linearized after 20 rad·s^−1^.

**Supplemental Figure 2:**
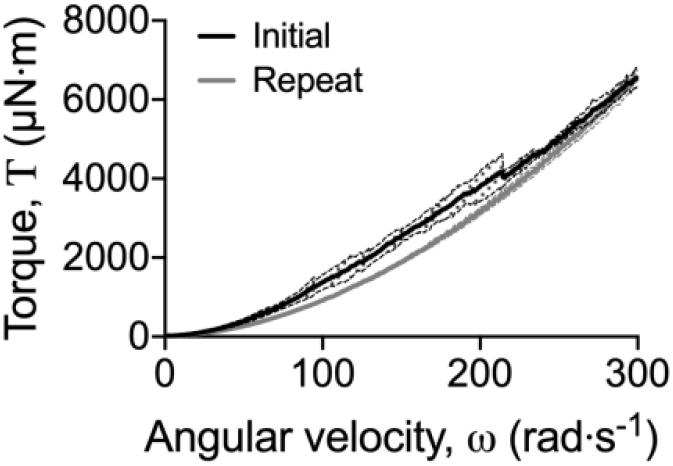
Repeated analysis does not lead to further biofilm removal. Untreated 5 day *S. gordonii* biofilms were analyzed by adapted rotating-disc rheometry. After the initial measurement (black) the assay was repeated (grey) to determine if remaining attached biofilm could be removed with subsequent analysis. Repeated analysis revealed no changes in torque, and the curve reached the same final point as the initial analysis. This indicates that no additional biofilm removal was detected with repeated analysis. Data presented as mean ± SD, N = 4.

**Supplemental Movie 1: Adapted rotating-disc rheometry measurement.** Left panel is a recording of the rheometry measurement for an untreated 5 d *S. gordonii* biofilm. Right panel indicates the corresponding torque – angular velocity data collection. Individual frames from the time lapse depicting separate biofilm detachment events are displayed in Fig 2. Time stamp is indicated in the top left hand corner (min: s). Playback rate is at 15 fps.

**Supplemental Movie 2: Transformed data collection.** Left panel is the same recording depicted in movie S1. Right panel depicts the corresponding torque – angular velocity data that has been linearized and transformed to emphasis the changes in torque. Individual frames from the time lapse depicting separate biofilm detachment events are displayed in Fig 2. Time stamp is indicated in the top left hand corner (min: s). Playback rate is at 15 fps.

